# A cortically gated sensory reference to the basal ganglia guides vocal reinforcement learning in songbirds

**DOI:** 10.64898/2026.11.11.765895

**Authors:** Aditi Agarwal, Constance Scharff, Daniela Vallentin

**Affiliations:** Max Planck Institute for Biological Intelligence, Seewiesen 82319, Germany; IMPRS Biological Intelligence, Seewiesen 82319, Germany; Graduate School of Systemic Neurosciences, Ludwig-Maximilians-Universität München, Planegg 82152, Germany; Freie Universität Berlin, Fabeckstraße 36a, Berlin 14195, Germany

**Keywords:** Birdsong, vocal learning, basal ganglia, Area X, premotor nucleus HVC

## Abstract

Songbirds learn to sing by copying a memorized tutor song ^1–4^. Reinforcement learning models of vocal imitation propose that a dopaminergic signal strengthens the cortico– basal ganglia synapses whose activity coincides with similarity to the tutor-like song ^5–7^. To do so, the circuit must access a representation of the tutor song. Whether such a reference is available within the songbird basal ganglia nucleus Area X, and how it arrives, is unknown. In juvenile male zebra finches that learn to imitate adult courtship songs, we found that lesions of Area X impaired imitation more severely the earlier they were made. Intracellular recordings from Area X in awake juvenile zebra finches showed that the tutor song evokes precise subthreshold depolarizations that inversely scaled with imitation proficiency throughout song learning. Silencing the upstream nucleus HVC reduced these tutor-evoked responses, whereas disinhibiting HVC enhanced them. An inhibitory gate in HVC thus supplies Area X with the temporally aligned tutor reference that reinforcement-driven vocal learning requires. These findings uncover a critical missing link in biological reinforcement learning, revealing how the brain structurally routes sensory templates to the basal ganglia to guide the acquisition of complex motor skills.

**Highlights:**

- Area X lesions in early development impair tutor-song imitation
- Tutor song evokes time-locked subthreshold responses in Area X neurons
- Depolarization inversely correlates with syllable similarity, allowing refinement
- HVC activity gates the strength of the tutor signal reaching Area X neurons

## Introduction

Learning to produce a complex motor sequence, whether human speech or the courtship song of a songbird, is thought to unfold in two stages: the learner first forms a sensory memory of a model and then uses that auditory template to guide the gradual refinement of its own vocal motor output ^8–10^. In the zebra finch, this process is developmentally gated. During an early sensitive period, a juvenile male listens to the song of an adult tutor. Over an overlapping and subsequent sensorimotor period, the juvenile practices tens of thousands of renditions, progressively molding his variable subsong toward a stereotyped copy of the tutor model ^1–4^. The tractability of this behavior, together with the constraint of a critical period of tutor song learning that closes around 90 days post hatch, has made the zebra finch an important model for exploring how a memorized target instructs vocal learning.

Central to the template and motor output comparison is the anterior forebrain pathway (AFP), a cortico-basal ganglia–thalamocortical loop. During the sensorimotor period, song-related premotor commands from the cortical nucleus HVC, which is critical for vocal learning and motor control in songbirds, reach Area X, a striatal nucleus that is specifically active during male juvenile song learning. A dedicated population of Area X-projecting neurons (HVC(X)) fire temporally precise, stereotyped bursts at specific moments during song production ^11,12^. Area X, in turn, is dominated by medium spiny neurons (MSNs) ^13,14^ that integrate time-locked cortical inputs from HVC and the Lateral Magnocellular Nucleus of the Anterior Nidopallium (LMAN) together with a dopaminergic projection from the ventral tegmental area (VTA) ^5,7,13,15–19^. A reinforcement learning framework has been proposed to explain how this circuit drives imitation. The VTA→Area X dopamine projection carries a reward-prediction-error-like signal that evaluates each rendition, increasing after better-than-expected performances and lacking after worse ones ^5,7,16,17,19^. By acting on an eligibility trace (a fading synaptic ‘memory’ for delayed reinforcement) at time-encoding HVC(X)→MSN synapses, this signal is proposed to selectively strengthen inputs that produce song variants resembling the tutor’s, thereby biasing premotor output toward the model. This bias is subsequently consolidated in downstream motor regions over a longer timescale ^6,20^. This framework is powerful, but it rests on an unstated prerequisite: to compute that a given variant is “more tutor-like”, the circuit must have access to a representation of the tutor song itself. Whether this representation is available within the AFP, and how it reaches Area X during learning remains unresolved.

Candidate loci for the tutor song memory have been identified largely outside the motor pathway. Following tutor exposure, sparse and highly selective neural responses to the tutor song emerge in higher-order auditory regions such as the caudomedial nidopallium (NCM) ^21^. While the NCM processes the initial perception of the tutor song, the nucleus Avalanche (Av) acts as a critical node within the evaluative circuitry, comparing the bird’s active sensorimotor feedback against this memorized template ^22^.

Within the motor pathway, the premotor nucleus HVC exhibits a performance-dependent representation of the tutor song ^23^ and is essential for encoding a sensory model that is capable of guiding vocal imitation ^24^. Furthermore, extracellular recordings of Area X neurons in anesthetized animals indicate that neurons within the AFP develop distinct song selectivity throughout the course of learning ^25,26^. However, it remains unclear whether Area X maintains an explicit representation of the tutor song that dynamically scales with imitation accuracy. If such a representation exists, the specific afferent populations providing this information are currently unknown. Resolving this gap is critical, as it constitutes the missing half of the current reinforcement-learning framework: elucidating not just how the Area X circuit utilizes an instructive signal but identifying where the sensory content of that signal ultimately originates.

Here we addressed these questions directly. Earlier work showed that Area X lesions disrupt adult song less when made later in development ^27^ but did not assess imitation fidelity to the tutor. To establish the time window during which Area X must be intact for faithful tutor song imitation, we lesioned it at several defined time points spanning the sensory and sensorimotor periods and quantified imitation fidelity. Next, to ask whether and how the tutor song is represented in Area X, we recorded responses to tutor-song playback across the critical period. Area X is predominantly populated by MSNs, whose low spike rates render subthreshold activity invisible to extracellular recordings. Therefore, we used intracellular recordings to resolve the membrane-potential dynamics evoked by the tutor song and track how these inputs change across developmental stages. Third, we identified the upstream source of this input using pharmacological manipulation. Together, these findings provide a circuit-level account of how information about the imitation similarity is established in Area X neurons to ensure reinforcement learning that drives vocal learning. By uncovering the missing neural link that supplies the sensory template, our results reveal a fundamental circuit mechanism by which the brain implements reinforcement learning to acquire complex behaviors.

## Results

### Area X lesions during the critical period impair tutor-song imitation in a developmentally graded manner

Lesion and ablation studies have long implicated Area X specifically in developmental learning ^27–30^. Interrupting the neural activity of Area X or HVC(X) projection neurons during the juvenile period disrupts normal song development. However, its effect on tutor imitation remains unresolved. To test whether and when Area X contributes to vocal imitation, we bilaterally lesioned Area X of juvenile male zebra finches at a series of ages during the critical period for song learning (46–81 days post hatch (dph), n = 8 birds). After each bird had been housed with a live adult tutor, we analyzed its song once it had crystallized (≥90 dph; Figure 1A). In birds lesioned early at 46-61 dph, although plastic song was present before and shortly after the lesion, the song failed to progress toward that of the adult, with the crystallized adult song bearing little resemblance to that of the tutor (Figure 1B, D). By contrast, birds lesioned later (74-81 dph) developed an adult song that closely recapitulated the tutor’s syllable structure but exhibited stuttering-like segments reported previously ^31^ (Figure 1C, E). We quantified copying across all lesioned birds using a maturity score computed from crystallized song (self-stereotypy × acoustic similarity to the tutor, normalized by tutor’s self-stereotypy). Maturity scores scaled positively with the age at which Area X was lesioned (Pearson correlation, r = 0.75, p = 0.0304; Figure 1F). Lesions earlier in the critical period yielded the most severely degraded, least tutor-like songs in adulthood whereas the copy fidelity improved progressively for lesions placed later in development. This finding is consistent with earlier reports that Area X lesions in the late stages of the critical period disrupted only those song elements that had not yet reached their adult form ^27^.

**Figure 1:**
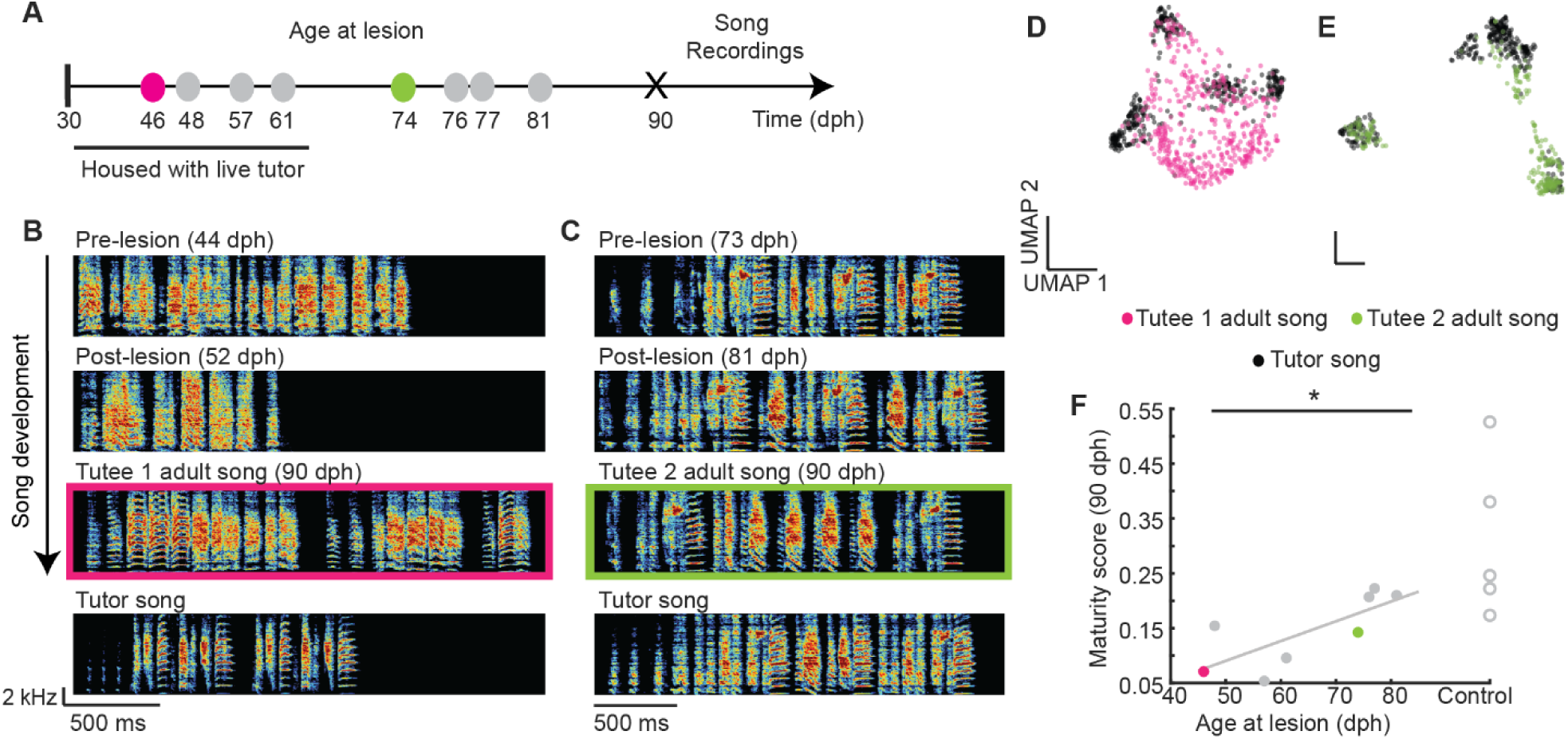
Area X lesions during the critical period impair tutor-song imitation in a developmentally graded manner. **A)** Time points when individual juveniles (n=8) received Area X lesions. Example birds are highlighted in pink (B) and green (C). **B)** Spectrograms of representative songs from a juvenile lesioned at 46 days post-hatch (dph), recorded at 44 dph (before the lesion), 52 dph (after the lesion), and in adulthood (>90 dph). **C)** Spectrograms of representative songs from a juvenile lesioned at 74 days post-hatch, recorded at 73 dph (before the lesion), 81 dph (after the lesion), and in adulthood (90 dph). **D)** and **E)** Latent embeddings of tutor song and adult tutee song (>90 dph) from birds in example B (pink) and C (green) visualized using UMAP. **F)** Maturity score (self-stereotypy x tutor imitation) of adult songs following Area X lesions (gray solid circles). Example birds from B and C are highlighted in pink and green, respectively (Pearson’s correlation, r = 0.75, p = 0.0304). Control birds (no lesioning) tutored with the same tutors are shown in gray open circles.

These results demonstrate that Area X is required for normal imitation of the male tutor song, and that this is developmentally graded. The earlier Area X is lesioned during the critical period, the more severely copying is impaired. This suggests that normally occurring neural dynamics within Area X progressively shape the male juvenile’s song toward the tutor model across the sensorimotor period.

**Figure S1:**
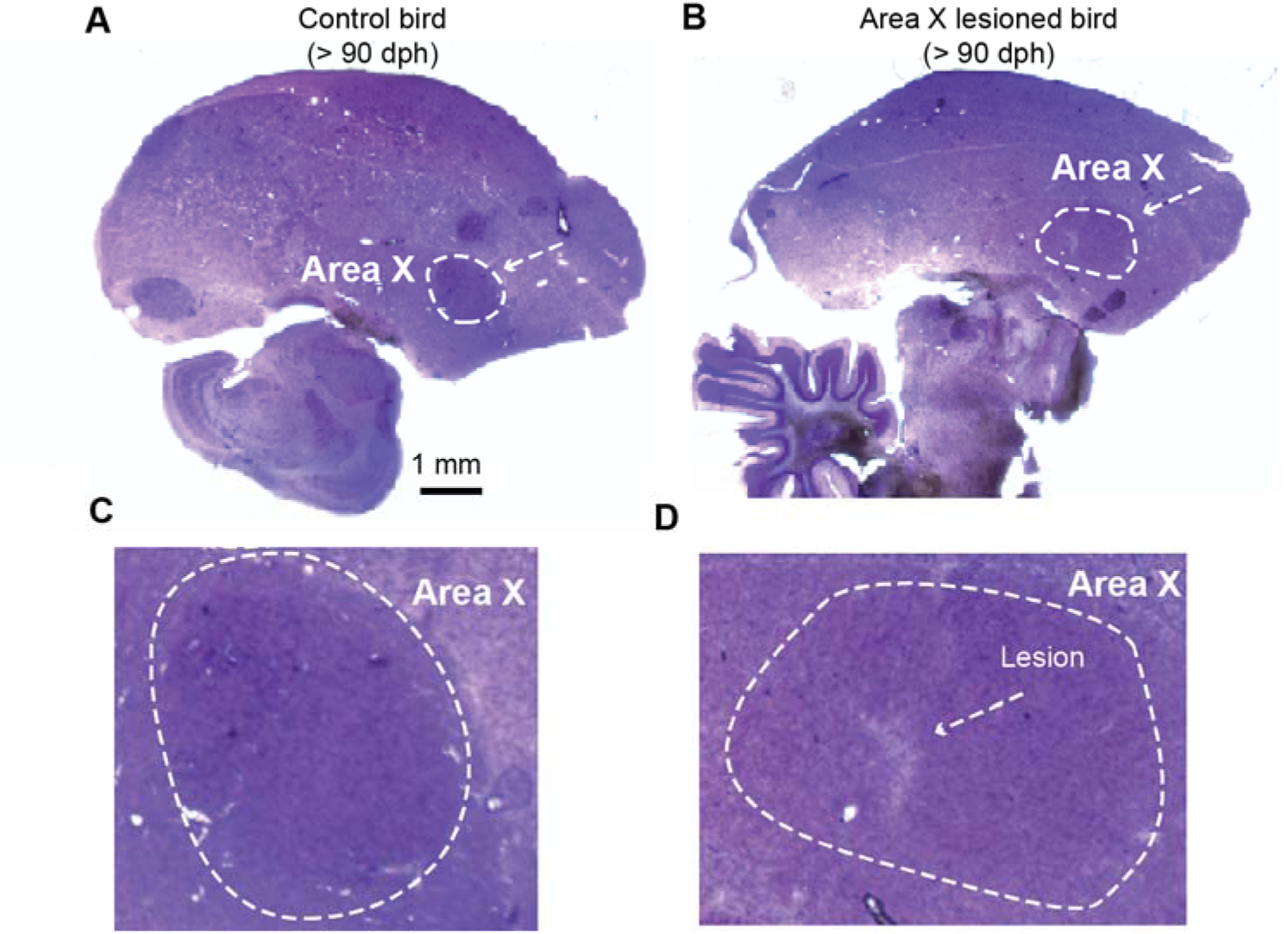
Area X histology in a 50 µm sagittal section stained with 1% w/v Cresyl Violet solution. Histology of adult brains (> 90 dph) of zebra finch with **A)** Control bird with an intact Area X. White arrow indicates Area X (stained with 1% w/v Cresyl Violet solution) **B)** Area X electrolytically lesioned at day 48 dph. White arrows indicate Area X. **C)** and **D)** Zoom-in of outlined Area X. White arrow points toward the lesion.

### Tutor song evokes spiking and subthreshold responses in Area X neurons

To dissociate the lesion’s interference with general AFP function from the contribution of Area X itself, we next asked whether Area X neurons represent the tutor song over the course of song learning. Previous studies using anesthetized juveniles showed that Area X neurons respond to song playback and develop selectivity for tutor song and the bird’s own song during learning ^25,32^. However, auditory responses in juvenile zebra finches can differ between awake, sleep and urethane anesthesia ^33,34^. This raises the possibility that responses measured under anesthesia may not fully reflect processing in awake birds, although this state difference has not been established specifically for Area X.

Area X is dominated by MSNs^13,14^ whose low spontaneous and evoked firing rates imply that much of the synaptic drive they receive remains subthreshold, and thus largely invisible to extracellular spike recording ^35^. We therefore performed sharp intracellular recordings from Area X neurons in awake juvenile zebra finches while playing back the song of their tutor, sampling neurons across the critical period (291 neurons from 19 birds; Figures 2A-C). Juveniles were housed with a live male tutor until ~63 dph (range: 55dph-73dph; Figure 2A), and recordings were distributed across ages spanning the sensorimotor period (38–82dph; Figure 2C). Unsupervised dimensionality reduction (UMAP) of the recorded spike waveforms, followed by HDBSCAN clustering, resolved seven neuronal types (Figure S2A) that differed systematically in their spontaneous firing rates (Figure S2B), ranging from sparsely firing, low-rate neurons, consistent with the MSNs that predominate in Area X, to higher-firing-rate cell types. This confirms that our recordings were not biased toward any single neuron class and that the tutor-evoked responses described below were sampled across the diversity of Area X neuron types, including the low-firing-rate neurons whose subthreshold input would be least accessible to extracellular methods. Further, upon visual examination, we did not see clustering based on recording age, suggesting that the intrinsic properties of Area X neurons remain consistent over the course of song development (Figure S2C).

**Figure 2:**
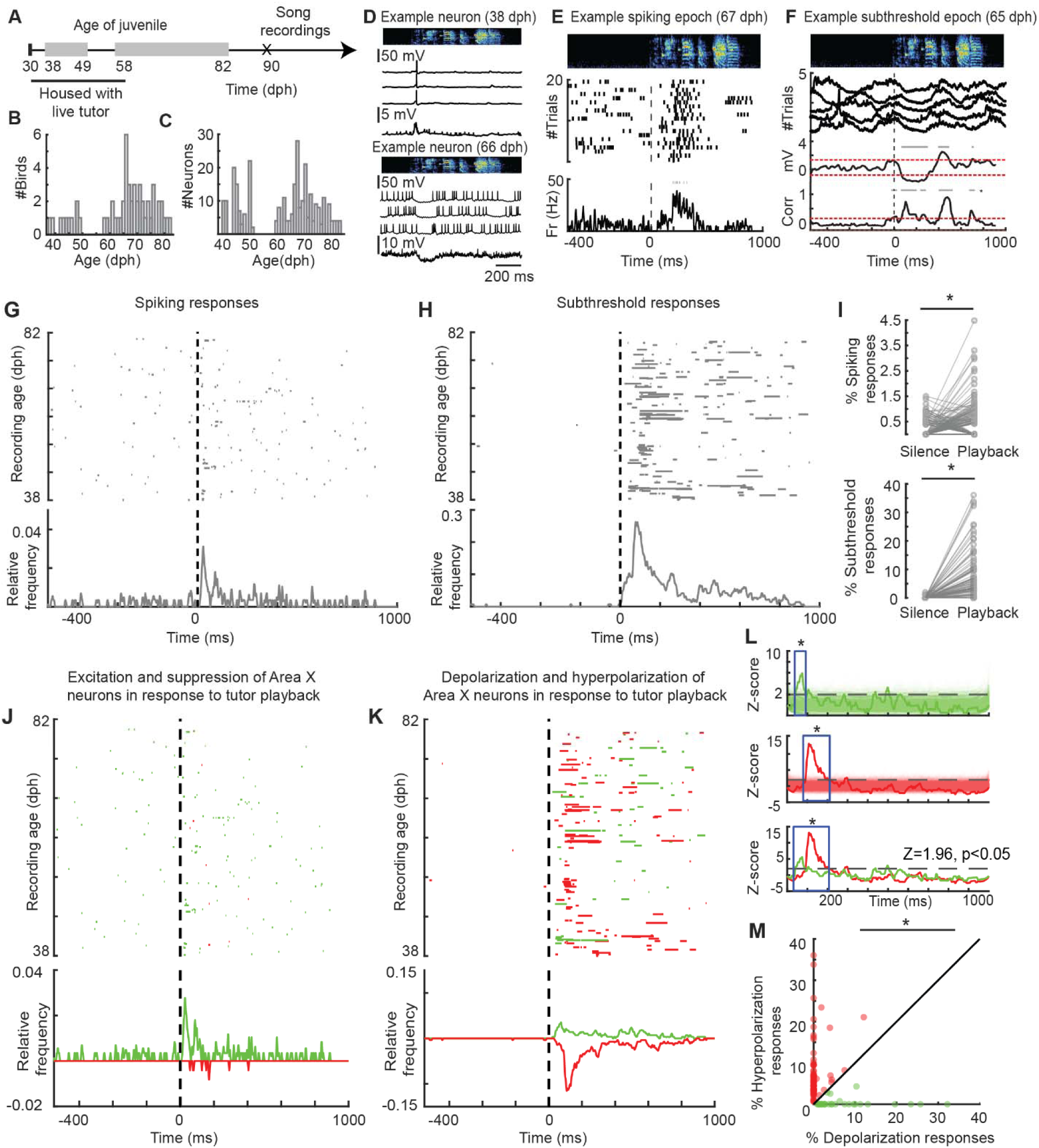
Tutor song evokes suprathreshold and subthreshold responses in Area X neurons of awake juvenile zebra finches. **A)** Juvenile zebra finches were housed with a live-male tutor until day 63 dph. Intracellular recordings were performed in awake juvenile zebra finches between 38-49 dph and 58-82 dph in response to playback of tutor song. **B and C)** Area X neurons in 19 juvenile birds (**B**) were recorded at different time points during the critical period (**C**). **D)** Example intracellular traces from neurons recorded in response to tutor song playback on day 38 (top) and day 66 (bottom) in two different birds. In each example, top panel depicts a spectrogram of the tutor’s song, middle panel shows three example traces from the respective neuron and at the bottom an average membrane potential trace across all trials. **E)** Dot spiking raster of example neuron with spiking responses recorded in a 67 dph old bird and **F**) example neuron with correlated subthreshold responses (bottom panel) recorded in a 65 dph old bird. Red dotted line indicates 99 % confidence interval in E and mean ± 3 x std in F (see Methods). Time epochs with a significant change in neural response during the playback are indicated by gray bars on top. **G)** Significant spiking response epochs from 291 neurons. Data pooled across all birds and recording sessions. Tutor song playback begins at time t = 0 ms. Each row represents one neuron. Bottom panel shows relative frequency of responses. **H)** Significant subthreshold response epochs across single neurons (top) and all neurons (bottom). **I)** Proportion of spiking (top, p = 0.0022, Wilcoxon signed-rank test) and subthreshold (bottom, p = 2.9e-29, Wilcoxon signed-rank test) responses during baseline versus playback of the tutor’s song. **J)** Excitation (green) and suppression (red) of spiking responses upon the playback of tutor song. **K)** Subthreshold responses color-coded by hyperpolarization (red) and depolarization (green) changes. **L)** (Top panel) Relative frequencies obtained from n = 1000 shuffles of depolarization subthreshold responses. Overlay: relative frequency from the actual distribution. Blue window indicates significant peak response to the playback of the tutor’s song (permutation test, n = 1000 shuffles, p < 0.05 (Z>1.96)). (Middle panel) Relative frequencies obtained from n = 1000 shuffles of hyperpolarization subthreshold responses. Overlay: relative frequency from the actual distribution. Blue window indicates significant peak response to the playback of the tutor’s song (permutation test, n = 1000 shuffles, p < 0.05). (Bottom panel) Overlay of relative frequencies obtained from real distribution of depolarization and hyperpolarization subthreshold responses as shown in L. **M)** Neurons that responded to tutor playback with either hyperpolarization or depolarization subthreshold change. Neurons with % hyperpolarization > % depolarization are represented in red. Neurons with % hyperpolarization < % depolarization are represented in green. Unity line represents equal duration of hyperpolarization and depolarization (p = 4.88e-06, Wilcoxon signed-rank test).

Individual Area X neurons responded to playback of the tutor’s song with membrane-potential modulation that was time-locked to the acoustic structure of the song (Figure 2D, Figure S3). At the population level, responsive neurons (n = 159/291 neurons) exhibited tutor-evoked spiking and subthreshold response epochs that were distributed throughout the playback (Figure 2E-F). These two non-exclusive response classes emerged after playback onset and persisted across the tutor motif and a brief window thereafter, rather than occurring at random times: neurons with elevations in firing rate (suprathreshold responses, n = 88 neurons, p = 0.0022, Wilcoxon signed-rank test; Figure 2G), and neurons with trial-reliable membrane-potential deflections (subthreshold responses, n = 109 neurons, p = 2.9e-29, Wilcoxon signed-rank test; Figure 2H). This indicates that the modulation of neuronal activity was driven by the tutor song and not by fluctuations in the awake bird’s baseline state.

Spiking responses comprised both excitation and suppression of firing rate (Figure 2I, J), and subthreshold responses comprised both depolarizing (activating) and hyperpolarizing (inhibitory) epochs (Figure 2I, K). Comparing each real distribution against shuffled data, both depolarizing and hyperpolarizing subthreshold responses exhibited a significant peak time-locked to the tutor playback (permutation test, n = 1000 shuffles, p < 0.05, Figure 2L), demonstrating that each polarity was stimulus-driven. We analyzed the timing of depolarizing and hyperpolarizing peak onset and found that depolarization (depolarization peak time = 72 ms) preceded hyperpolarizations (hyperpolarization peak time = 109 ms) (Figure 2L, bottom panel), indicating an external excitatory signal directly recruits local inhibition. Across birds, tutor song elicited both hyperpolarizations and depolarizations in a small subset of neurons (15/109 responsive neurons). The proportion of hyperpolarizing epochs (n = 70 epochs in 109 responsive neurons) exceeded that of depolarizing epochs (n = 24 epochs in 109 responsive neurons, p = 4.88e-06, Wilcoxon signed-rank test, Figure 2M). Thus, while tutor playback elicited both depolarized and hyperpolarized Area X neurons, the net subthreshold drive was biased toward inhibition.

Together, these recordings demonstrate that Area X neuron activity in awake juvenile zebra finches represents the tutor song throughout the critical period and that this representation is expressed to a substantial degree at the subthreshold level.

**Figure S2:**
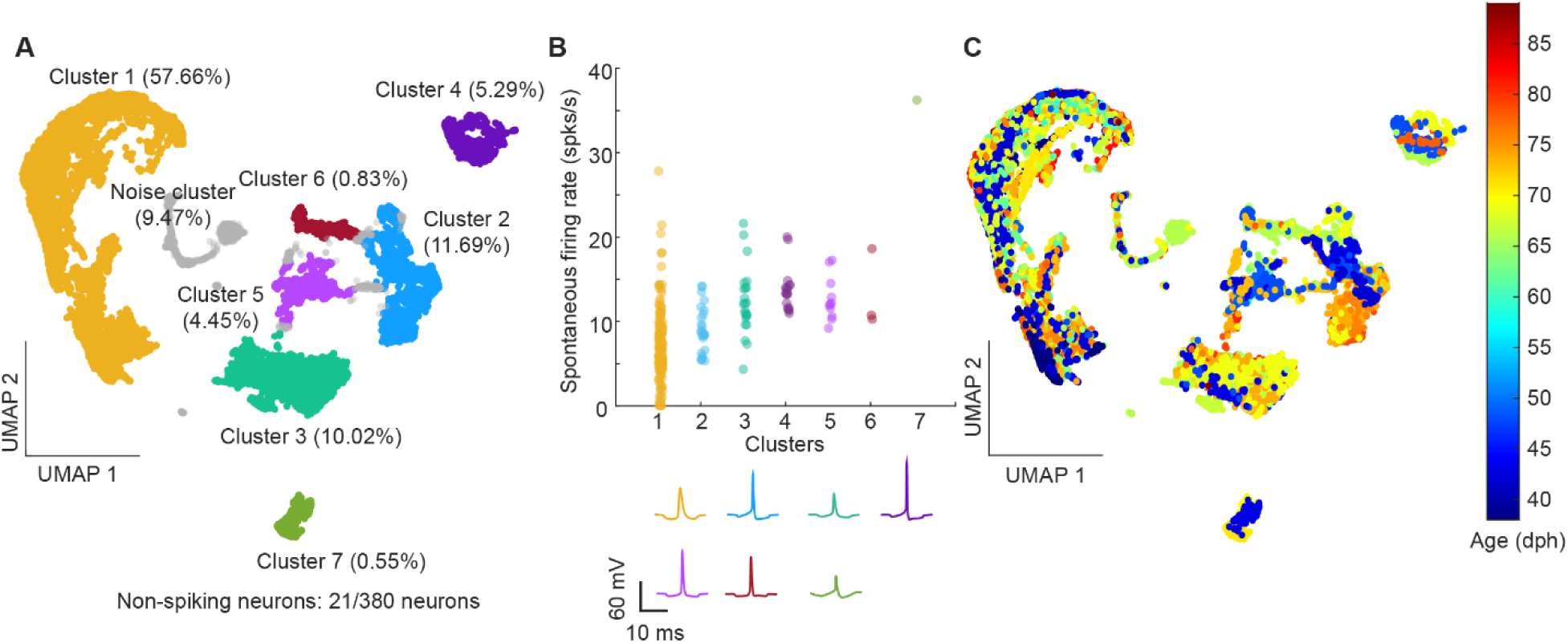
Spike waveform analysis reveals different neuron clusters. **A)** Spike-waveform reduction and visualization using UMAP reveals seven clusters of neurons. If 80% of the spikes from a neuron fell into the same cluster, it was taken to be a valid neuron otherwise labelled as noise (shown in gray) **B)** Cluster-wise spontaneous firing rates of neurons. (Bottom) Median spike waveform from respective clusters. **C)** Spike waveform latents color-coded by age of recording.

**Figure S3:**
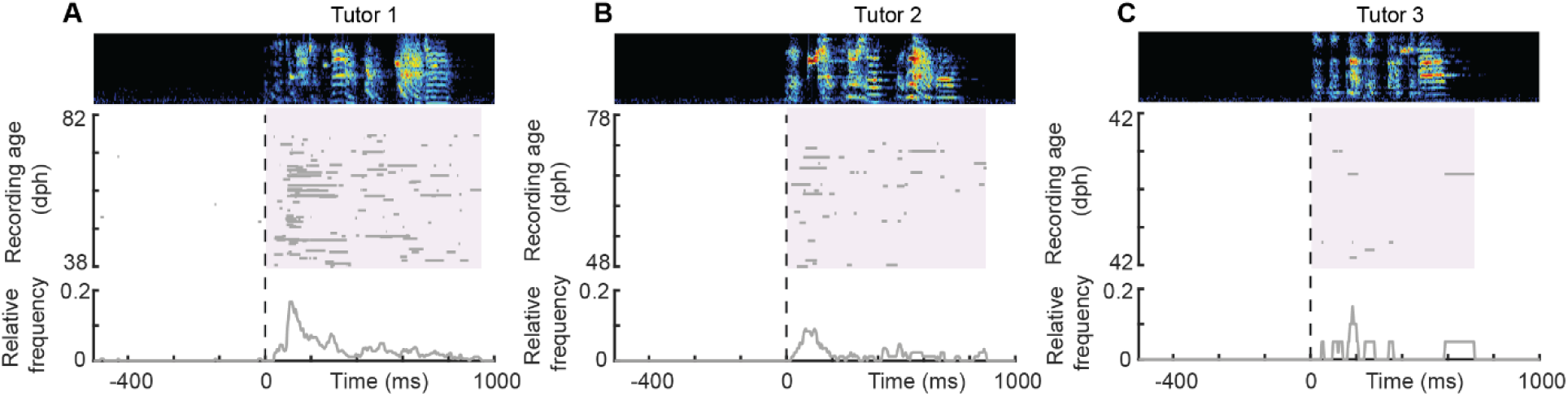
Subthreshold responses to tutor song playback in tutees over the course of song learning: **(A-C)** Top panel: spectrogram of the respective tutor song. Middle panel: subthreshold response epochs aligned to the onset of the tutor song playback in the order of increasing age for all tutees tutored with the respective tutor. Each row represents one neuron. Shaded region represents the average duration of tutor song calculated from all playbacks with an additional 100 ms (tutor 1: 933.74 ± 30.34 ms, tutor 2: 870.83± 5.27 ms, tutor 3: 715.40 ± 1.50 ms; mean ± std). Bottom panel: Relative frequency of subthreshold response epochs aligned to the playback of the tutor song.

### Depolarizing responses in Area X scale inversely with tutor-song imitation

Next, we asked whether the tutor song representation in Area X changes over the course of the critical period. If tutor-evoked subthreshold activity of Area X neurons helps guide imitation, its magnitude should track how much of each element has been learned. We therefore compared response magnitude for individual syllables against each syllable’s current imitation proficiency. Because syllables within a single bird’s song, recorded at the same age, are typically imitated to different degrees, this per-syllable comparison dissociates a learning-related effect from the bird’s chronological age (Figure 3A-C).

**Figure 3:**
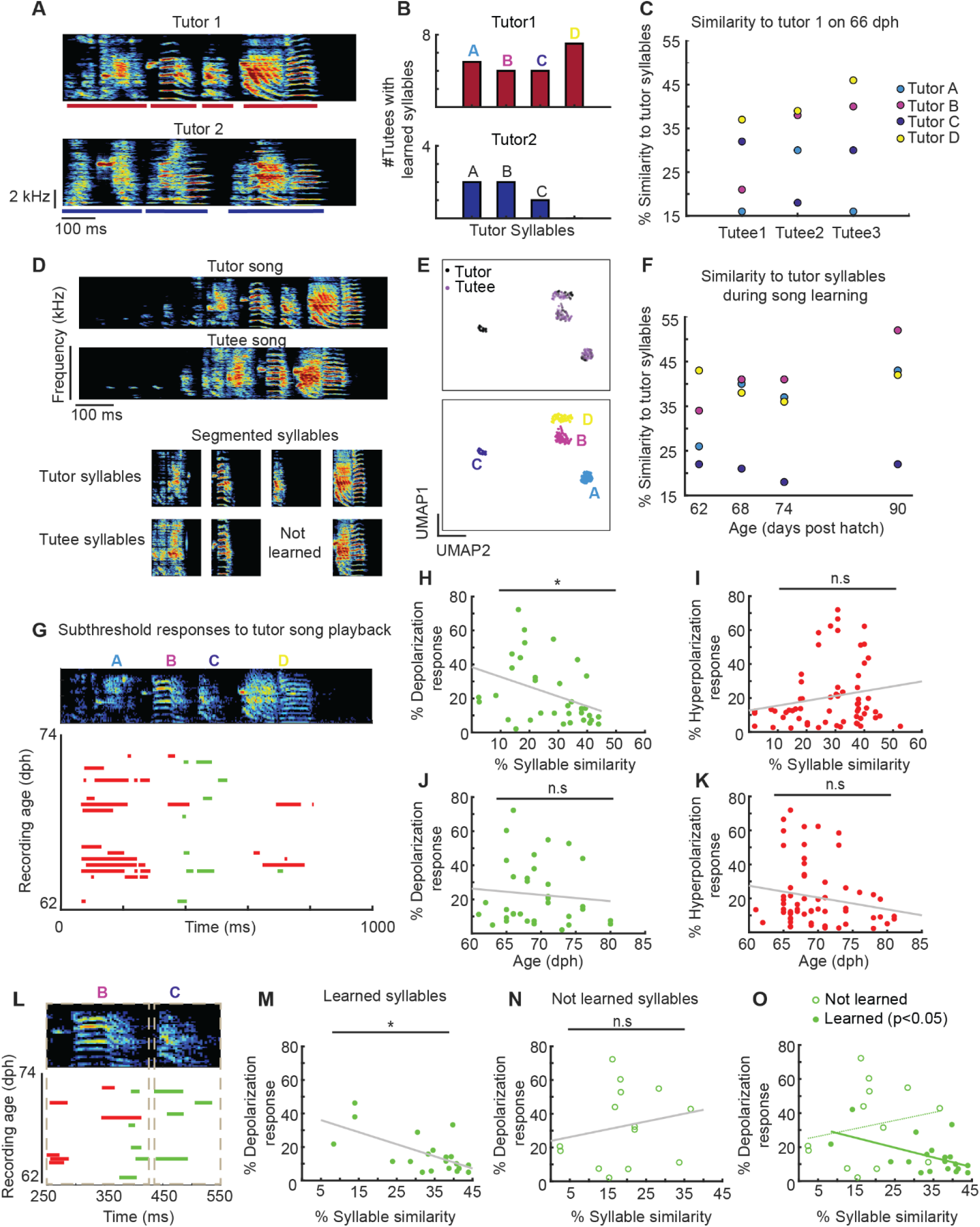
Depolarization response epochs in Area X correlate with proficiency in tutor song syllables. **A)** Representative song spectrograms for tutors used for live-tutoring (n= 12 juveniles individually tutored). Individual syllables are marked with a bar under each syllable. Tutor 1 had four syllables and tutor 2 had three syllables **B**) Different syllables learned by tutees from their corresponding tutors at the end of the recording period (90 dph) **C**) Similarity scores for tutor 1 syllables-A,B,C,D for different tutees on 66 dph. Every point corresponds to a similarity score between the tutor and corresponding tutee syllable (see Methods). **D**) Example tutor and tutee song (>90 dph). (Bottom panel) Segmented tutor and tutee syllables **E)** (Top panel) 2D UMAP visualization of latents obtained by reducing all tutor and tutee (adult song >90 dph) motif syllables. (Bottom panel) Syllable clusters obtained from HDBSCAN clustering on latent embeddings from E (top). Syllables ‘A’,’B’,’D’ from the tutor are incorporated into the crystallized repertoire of tutee (learned syllable = overlap with tutor syllable in the latent space), syllable ‘C’ is not learned **F)** %Similarity of tutee (from Figure 3D) with corresponding tutor syllables over the course of song learning. **G**) (Top panel) Spectrogram of tutor song. (Bottom panel) Subthreshold responses of all neurons (depolarization shown in green, hyperpolarization shown in red) recorded between day 62 and 74 upon playback of tutor’s song (n = 26 neurons) in example juvenile shown in **D**. **H)** Depolarization subthreshold responses normalized by the duration of the syllable plotted against similarity between tutor syllable and corresponding tutee syllable measured on the day of respective neural recording (F(1,31) = 6.3181, p = 0.017, linear mixed-effect model). **I)** Hyperpolarization subthreshold responses normalized by the duration of the syllable plotted against similarity between tutor syllable and corresponding tutee syllable (F(1,59) = 0.58719, p = 0.45, linear mixed-effect models). **J)** Depolarization subthreshold responses normalized by the duration of the syllable plotted against age of the bird (F(1,31) = 0.60, p = 0.443, linear mixed-effect model). **K)** Hyperpolarization subthreshold responses normalized by the duration of the syllable plotted against age of the bird (F(1,59) = 1.2809, p = 0.26232). **L**) Zoom in from figure G: Top panel: Copied syllable ‘B’ (overlap in latent space with tutee syllable) vs un-incorporated syllable ‘C’ (no-overlap in the latent space) Bottom panel: depolarization and hyperpolarization subthreshold responses shown in green and red respectively. Dotted boxes represent syllable boundaries with an additional 30 ms. %Subthreshold responses for copied syllables (**M**) not learned syllables (**N**) **O**) Depolarization responses inversely scale with imitation proficiency for learned syllables (F (1,29) = 4.5843, p = 0.0408, linear mixed-effect model) but not for unlearned syllables (F(1,29) = 0.578, p = 0.453, linear mixed-effect model).

To objectively classify syllables as ultimately learned versus unlearned, we segmented both tutor and tutee songs into syllables and embedded all tutor and adult tutee (>90 dph) motif syllables in a common low-dimensional space (UMAP), clustering them with HDBSCAN (Figures 3D, E). Because zebra finches can learn song structure at the subsyllabic level, we used an objective classifier to assess tutor–pupil syllable correspondences ^36^. A tutee syllable was scored as a learned copy of a tutor syllable when the two overlapped in this latent space, otherwise it was scored as not learned. To quantify imitation success syllable-by-syllable, syllables produced on a given day (current proficiency) were correlated with corresponding tutor syllables based on whether or not they were eventually learned. For ‘not yet learned’ tutor syllables, a similarity score was assigned by averaging similarity of this tutor syllable with all other tutee syllables produced that day (Figure 3D-F). Across tutees, individual tutor syllables were acquired with differing frequency and fidelity (Figures 3B-C), providing a graded, syllable-resolved index of imitation proficiency.

We then related, for each syllable, the tutor-evoked subthreshold response (the proportion of the syllable when a significant subthreshold response in the neural activity was observed, normalized by syllable duration) to the syllable’s similarity to tutor (Figure 3G-K). We limited our analysis to Area X neurons recorded between 60-82 dph (n=188 neurons, n=12 juveniles) since syllable identities were not consistently possible for birds younger than 60 dph in our dataset. The magnitude of depolarizing responses was negatively correlated with imitation proficiency i.e. well-copied syllables evoked smaller responses and poorly copied syllables evoked larger depolarizations (Figure 3H, depolarization predicted by proficiency: F(1,31) = 6.3181, p = 0.017, linear mixed-effect model, n = 25/188 neurons with 34 responses in 10 birds). Hyperpolarizing responses, however, showed no relationship to syllable similarity (Figure 3I, Hyperpolarization predicted by proficiency: F(1,59) = 0.58719, p = 0.45, linear mixed-effect models, n = 62 responses in 12 juveniles). We did not find a correlation between subthreshold neural responses and age of the juvenile (depolarization predicted by age: F(1,31) = 0.60, p = 0.443, Figure 3J; hyperpolarization predicted by age: F(1,59) = 1.2809, p = 0.26232, Figure 3 K; linear mixed-effect model).

We next asked whether the putative teaching signal expressed as excitatory depolarizing input depends on whether a tutor element is being refined or potentially incorporated into the juvenile’s song. We predicted that, for incorporated syllables, depolarization would vary with ongoing imitation proficiency, decreasing as the juvenile’s copy became more accurate. For syllables that had not been incorporated, we predicted that depolarization would be present during playback of the corresponding tutor syllable but would not vary with the proficiency score (because they are not being refined).

Consistent with this prediction, depolarization magnitude covaried with proficiency for incorporated (learned) syllables (F (1,29) = 4.5843, p = 0.0408, linear mixed-effect model, n = 20 responses in 8 juveniles; Figure 3L-O), whereas we detected no association between depolarization magnitude and proficiency for un-incorporated (‘not learned’) syllables (F(1,29) = 0.578, p = 0.453, linear mixed-effect model, n =14 responses in 8 juveniles; Figure 3L-O). Together, these findings suggest that depolarization relates to ongoing refinement for incorporated syllables, while also being elicited by tutor syllables that have not yet been incorporated. Therefore, the prominent hyperpolarizing (inhibitory) input identified above (Figure 2M) is present regardless, whereas the depolarizing (excitatory) input is graded by learning, largest for the elements the bird has yet to master and diminished as imitation improves (depolarizing input for learned syllables: 11.51%, (IQR: 7.25,19.92); not learned syllables: 31.48%, (IQR: 11.17,52.79); median depolarization response, IQR (25-75); p = 0.0342; Wilcoxon ranksum test).

To assess whether the Area X subthreshold neural responses were specifically representing the memorized tutor song or also the bird’s own song (BOS), we analyzed a subset of neurons from our dataset (n=91 neurons) which received playback of both BOS and the tutor song (Figure S4). BOS provides a comparison stimulus because it contains syllables that can resemble those of the tutor song, although the syllables may differ in their acoustic similarity and sequence position. During tutor song playback, the subthreshold depolarization associated with each tutor syllable was negatively related to the similarity between that tutor syllable and the corresponding juvenile syllable produced on the same day (F(1,19) = 15.3824; p = 0.00092, linear mixed-effect model, n = 21 depolarization epochs, Figure S4D). In contrast, during BOS playback, Area X neural responses were not detectably related to the similarity of the corresponding BOS elements to the tutor syllables (F(1,19) = 0.2202; p = 0.6443, linear mixed-effect model, n = 13 depolarization epochs, Figure S4 B-D). The response–similarity slopes differed between playback conditions (Playback × Similarity interaction: F(1,19) = 6.5523; p = 0.01916). This pattern is consistent with a tutor song specific association between neural response and the bird’s current imitation of individual tutor syllables, rather than a general association between neural responsiveness and tutor similarity.

Together, these results indicate that the tutor song representation in Area X is sculpted by the state of learning. Depolarizing responses are strongest for the syllables a bird has yet to master and wane as its copy improves, exactly the profile of a signal reporting how much of the tutor model is acquired. This depolarizing input is therefore the component most plausibly instructing song refinement, and its decline with proficiency predicts a learning dependent decline over developmental time, linking it to the developmental contribution of Area X revealed by our lesions.

**Figure S4:**
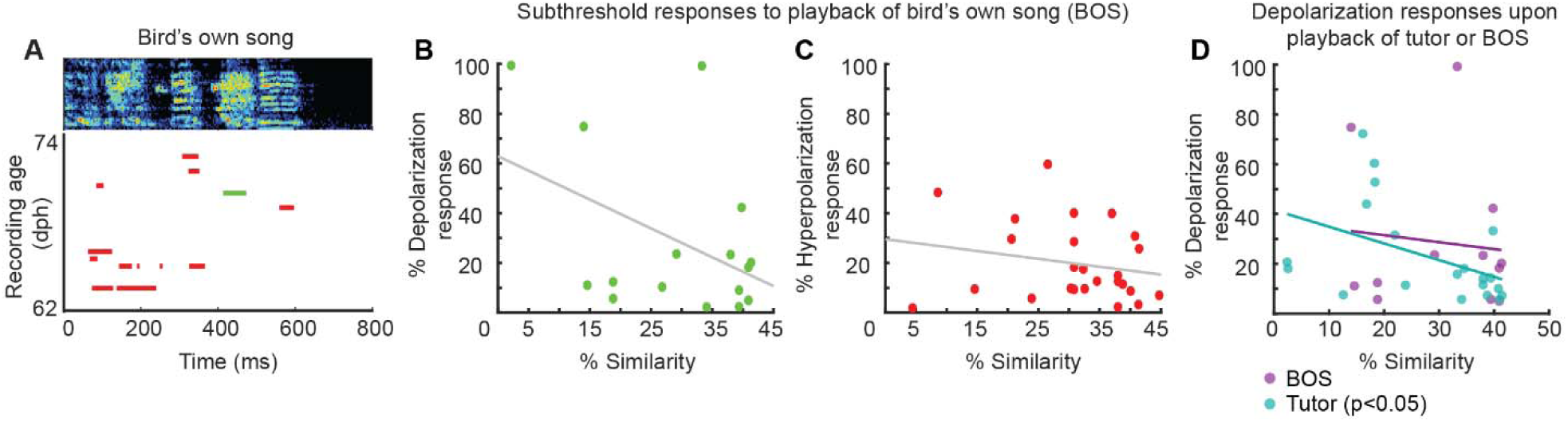
Area X responses track syllable-specific imitation during tutor-song playback, but not BOS playback. **A)** Responses to BOS (n = 23 neurons recorded between 62-74 dph) in the example bird shown in Figure 3G. **B)** Depolarization subthreshold responses normalized by the duration of the syllable plotted against similarity between the tutee syllable and its corresponding tutor syllable measured on the day of respective neural recording (n = 100 neurons, n = 9 birds, n = 16 response epochs). **C)** Hyperpolarization subthreshold responses normalized by the duration of the syllable plotted against similarity between the tutee syllable and its corresponding tutor syllable measured on the day of respective neural recording (n = 100 neurons, n = 9 birds, n = 25 response epochs) **D)** % Depolarization responses upon the playback of tutor song (blue) and birds own song (purple) in a subset of neurons (n = 91 neurons) plotted against the similarity to either tutor syllables (for BOS playback) or respective tutee syllables (for tutor song playback) on the day of recording (linear mixed effects model, tutor: F(1,19) = 15.3824; p = 0.00092, n=21 depolarization epochs; BOS: F(1,19) = 0.2202; p = 0.6443, n = 13 depolarization epochs; interaction (playback x similarity): F(1,19) = 6.5523; p = 0.01916).

### An inhibitory gate in HVC controls the tutor song representation in Area X

Where does this instructive signal originate? Two observations point upstream to HVC. First, the depolarizing and hyperpolarizing components of the Area X response are temporally distinct (Figure 2K–M), suggesting that excitation and inhibition arrive by separate routes, with the excitatory drive consistent with the glutamatergic HVC(X) projection that constitutes Area X’s principal cortical input. Second, inhibition within HVC has been shown to gate auditory-related activity in a learning-dependent manner, strengthening around syllables as they are acquired ^23^. We therefore hypothesized that the tutor-evoked, instructive signal in Area X originates in HVC and is set by an inhibitory gate within HVC.

To test whether HVC is the source of the tutor-evoked input to Area X, and whether that input is controlled by inhibition within HVC, we manipulated HVC pharmacologically while making simultaneous intracellular recordings from Area X neurons in age-matched juveniles (61-89 dph) during tutor song playback (Figure 4). We used two manipulations with opposite predicted effects on the HVC “gate”: muscimol, a GABA_A_ receptor agonist, to suppress HVC output and close the gate; and gabazine, a GABA_A_ receptor antagonist, to block intra-HVC inhibition exerted by HVC interneurons, thereby opening the gate by activating HVC(X) projection neurons (Figure 4A-C).

**Figure 4:**
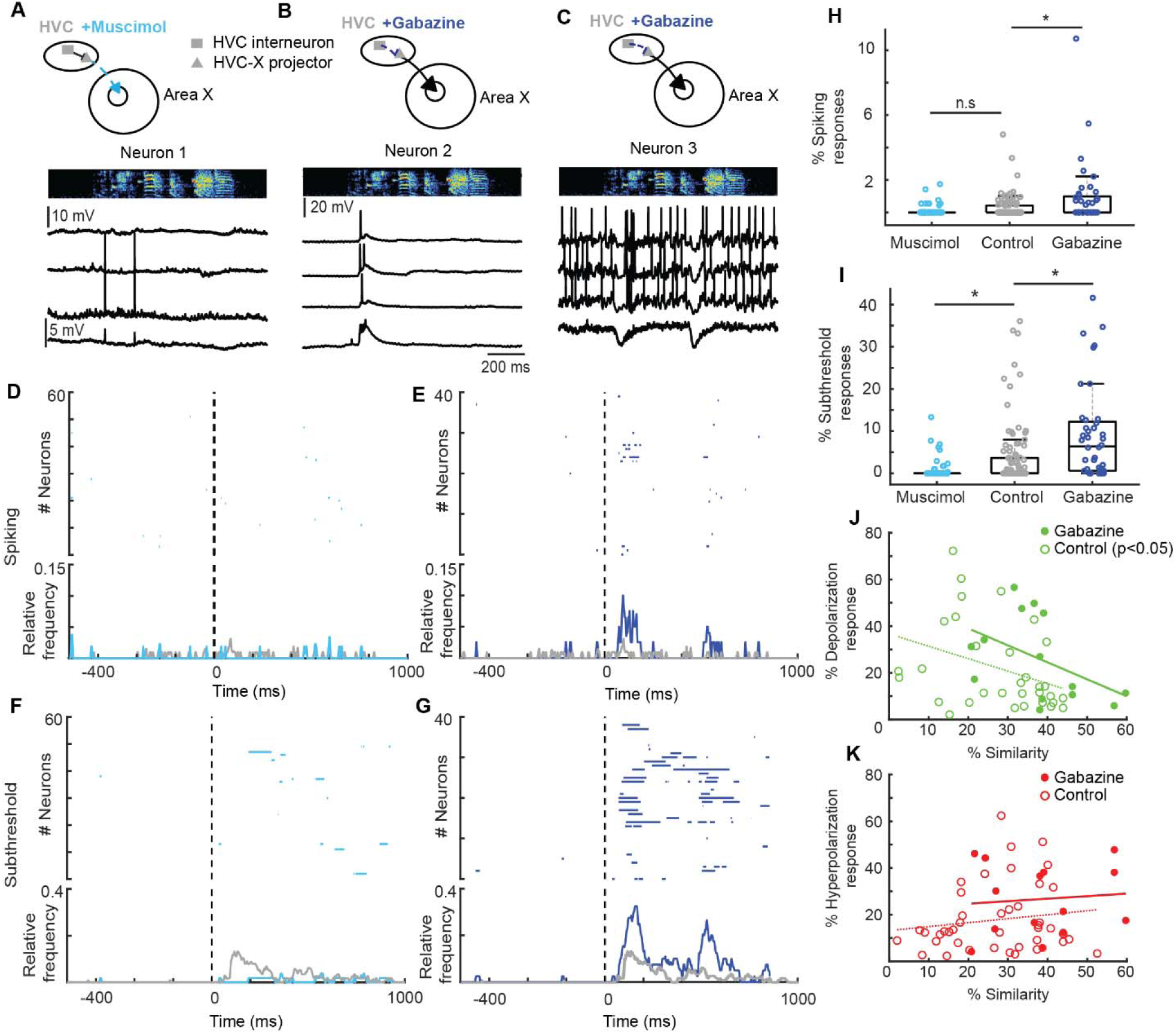
An inhibitory gate in HVC controls the tutor-song representation in Area X. **A)** Example neuron recorded in Area X during tutor song playback while muscimol was infused onto HVC. **B, C)** Example neurons recorded in Area X in response to gabazine in HVC. **D and E)** Spiking profiles of all neurons recorded in Area X in response to HVC manipulation. (Bottom panel) Relative frequencies of all neurons recorded compared against age-matched control birds tutored with the same tutor (in gray). **F and G)** Subthreshold profiles of all neurons recorded in Area X in response to HVC manipulation. (Bottom panel) Relative frequencies of all neurons recorded compared against age-matched control birds (in gray). **H and I)** Proportion of spiking (H) and subthreshold (I) responses across all neurons recorded in Area X in response to HVC manipulation compared to control birds. **J)** % Depolarization response in Area X neurons with (solid circles and solid line) and without gabazine in HVC (open circles with dashed line) per tutor syllable plotted against its similarity to corresponding tutee syllable (without gabazine: F(1,52) = 5.1006, p = 0.02814; with gabazine: F(1,52) = 1.2077,p = 0.27686, linear mixed-effect model). **K)** % Hyperpolarization response in Area X neurons with (solid circles and solid line) and without gabazine in HVC (open circles with dashed line) per tutor syllable plotted against its similarity to corresponding tutee syllable (without gabazine in HVC: F(1,88) = 0.58424, p = 0.4467; with gabazine in HVC: F(1,88) = 0.4171, p = 0.52007, linear mixed-effect model).

Silencing HVC output with muscimol did not change the spiking responses to tutor song in Area X (playback (control: 0, (IQR: 0–0.42%), n = 113 neurons; muscimol: 0 (IQR: 0-0%), n = 53 neurons; (median, IQR (25-75)); p = 0.1775, Wilcoxon ranksum test, Figure 4D, H). However, this manipulation reduced tutor-song-evoked subthreshold activity in Area X neurons during playback (control: 0, (IQR: 0–3.63%); muscimol: 0, (IQR: 0-0%); (median response, IQR (25-75)); p = 0.0091, Wilcoxon ranksum test, Figure 4F, I). Therefore, with limited HVC input the tutor song no longer drove Area X subthreshold activity, demonstrating that HVC is required for the representation rather than merely modulating it.

Conversely, removing the impact of inhibition within HVC with gabazine opened the auditory gate and enhanced tutor-evoked responses in Area X. Disinhibiting HVC(X) projection neurons increased both the spiking responses (control: 0, (IQR: 0–0.42%), n = 113 neurons; gabazine: 0, (IQR: 0-0.98%), n = 40 neurons; (median response, IQR (25-75)); Wilcoxon ranksum test, p = 0.0060, Figure 4B, E, H) and the subthreshold responses (control: 0, (IQR: 0–3.63%); gabazine: 6.41, (IQR: 0.58,12.24%), n = 40 neurons; (median response, IQR (25-75)); p = 1.40e-05, Wilcoxon ranksum test, Figure 4C, G, I) of Area X neurons to tutor playback relative to control, indicating that intra-HVC inhibition normally constrains how strongly the tutor song drives Area X. Disinhibiting HVC(X) neurons increased Area X spiking during tutor-song playback, but did not produce a detectable increase in baseline firing. Baseline firing rates did not differ between age-matched controls and the gabazine condition (Wilcoxon rank-sum test, p = 0.5938; Fig. S5A). Spiking therefore remained time-locked to tutor song playback, rather than reflecting a measurable increase in spontaneous activity. Overall, the pharmacological manipulation affected song production only transiently, and song had fully recovered within two days (Fig. S5B, C).

We hypothesized that inhibitory interneurons in HVC gate activity in HVC(X) neurons, thereby regulating the tutor-related relationship between imitation proficiency and Area X responses. Accordingly, reducing inhibition in HVC with gabazine should weaken the association between proficiency and subthreshold depolarization responses. Consistent with this prediction, HVC disinhibition altered the proficiency–depolarization relationship (control: F(1,52) = 5.1006, p= 0.02814, gabazine: F(1,52) = 1.2077, p = 0.27686, linear mixed-effect model, Figure 4 J). By contrast, we found no evidence that HVC disinhibition altered the hyperpolarization responses of Area X neurons (control: F(1,88) = 0.58424, p = 0.4467; Gabazine: F(1,88) = 0.4171, p = 0.52007, linear mixed-effect model, Figure 4 K).

Together, these bidirectional manipulations establish HVC as the source of the tutor-song-evoked input to Area X and confirms inhibition within HVC as the gate that sets its strength ^23,37^. This provides a mechanism for the learning-dependent decline in depolarizing responses that we observed (Figure 3). As a syllable is acquired, inhibition in HVC strengthens ^23^, progressively closing the gate and reducing the tutor-evoked drive transmitted to Area X. HVC therefore supplies Area X with a representation of the tutor song whose magnitude is continuously scaled by an inhibitory gate that tracks the state of learning, furnishing the instructive input that reinforcement learning of song imitation requires, and closing the loop between the memorized tutor reference and the AFP circuitry that acts upon it.

**Figure S5:**
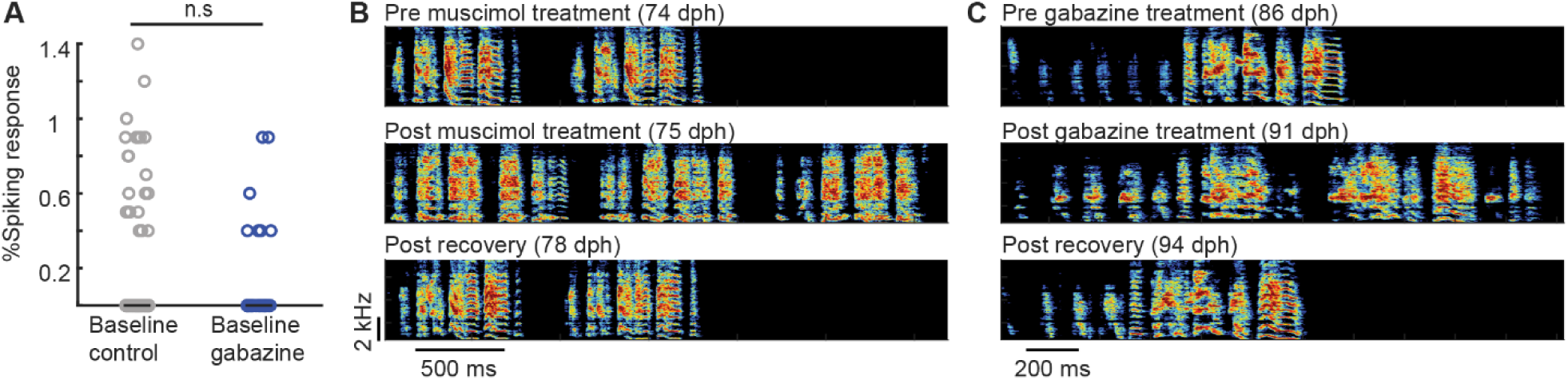
The Impact of Muscimol and Gabazine Treatments on Song and Baseline Neural Activity. **(A)** Comparison between spiking response during baseline control and baseline gabazine (Wilcoxon rank-sum test, p = 0.5938) **(B)** Spectrograms of example songs pre-muscimol treatment at 74 days post-hatch (dph), post-muscimol treatment at 75 dph, and post-recovery at 78 dph. **(C)** Spectrograms of example songs pre-gabazine treatment at 86 dph, post-gabazine treatment at 91 dph, and post-recovery at 94 dph.

## Discussion

Reinforcement learning models of vocal imitation require that the song circuit accesses a representation of the tutor song to evaluate which variants are more tutor-like. Here, we show that such a reference is available in Area X (Figure 1 and 2) and that it takes the form of a learning-graded subthreshold depolarizing signal (Figure 3). Because depolarizing responses were strongest for syllables the bird had yet to copy and weakened as imitation improved, they appear to track the residual gap between the bird’s song and the tutor model, a signal of how much remains to be acquired. Our manipulations indicate that this dynamic is imposed within HVC. Silencing HVC abolished tutor-evoked responses in Area X, whereas relieving intra-HVC inhibition of HVC(X) neurons enhanced them. Together with the finding that HVC inhibition strengthens around syllables as they are learned ^23^, this suggests that the gate progressively closes on mastered elements while remaining open for what is still being learned, concentrating the instructive drive where it is most needed (Figure 4). These findings reveal the crucial missing element of the reinforcement learning framework: the mechanism by which an internal model is used to refine imitation. Ultimately, this reveals a fundamental biological mechanism for reinforcement learning, demonstrating how the brain evaluates motor actions against memorized goals to master complex, learned behaviors.

Our results reveal how the target memory is delivered to the AFP circuit to guide evaluation and learning. During natural, self-directed practice, dopamine release in Area X rises for renditions that are closer to the song’s crystalized adult form and decreases after renditions that are farther from it, tracking performance quality on a rendition-by-rendition basis ^5,7^. However, this does not establish whether the neural evaluation signal coming via dopaminergic inputs specifically uses the tutor song as its comparison target. For a rendition to be evaluated as more or less tutor-like, the circuit must have access to a tutor-derived reference. Our data identify where that reference enters AFP the circuit, an HVC-gated input to Area X whose influence varies not only with the current performance proficiency but also the status of incorporation, thereby demonstrating copying from a memorized target. This interpretation, however, assumes that when juveniles hear the tutor, the brain combines that input with information about their recent singing. Whether such performance-related information is retained as a memory trace, and where it might be represented, remain areas for future research.

Notably, dopamine release in Area X is driven not only by midbrain dopamine neurons but also by song premotor input acting through local cholinergic signaling ^5^, thereby suggesting that HVC influences Area X through multiple channels. The inhibitory gate we describe may therefore shape both the direct glutamatergic tutor signal and, indirectly, the premotor drive onto the dopaminergic system. Additionally, mapping waveform-defined cell types and anatomically and transcriptionally distinct neurons would further clarify the inputs that determine imitation vocal learning.

Synapse-specific optogenetic and chemogenetic manipulations recently identified the HVC-to-Area X MSN connection in the songbird basal ganglia as driving the acquisition and expression of rapid, learning-related vocal changes ^6^, the plastic node posited by reinforcement learning models of imitation. Our findings show that this same connection conveys a gated, tutor-defined instructive signal, while the dopamine studies show that its plasticity is under reinforcement control. A coherent picture thus emerges in which this HVC→Area X synapse integrates a temporally structured premotor eligibility signal, and a gated representation of the tutor target that the reinforcement circuitry needs in order to be meaningful.

The observation of brief subthreshold depolarizations during specific elements of the tutor’s song suggests that neuronal activity relevant to learning may extend beyond suprathreshold firing and could provide a temporally precise signal for synaptic modification. In other systems, brief subthreshold depolarizations have been shown to function as Hebbian signals that are capable of driving long-term plasticity ^38^. Although we did not directly test whether the events observed here induce synaptic modifications, their occurrence during behavior is consistent with the possibility that they contribute to experience-dependent circuit refinement.

Beyond the avian song system, these findings outline a conserved circuit logic for biological reinforcement learning. Evaluating ongoing motor output against a memorized sensory template is fundamental to developmental skill acquisition, including human speech. Avian and mammalian cortico-basal ganglia networks share deep homology ^20^. We propose that these analogous projections route sensory reference templates to the basal ganglia to evaluate developmental motor “babbling.” Broadly, this establishes a conserved vertebrate mechanism by which sensory memory guides motor plasticity to master complex behaviors.

### Limitations of the study

Our pharmacological manipulations of HVC activity affected all outputs rather than the tutor-related component selectively. In turn, the responses in Area X neurons were measured to playback rather than during singing, leaving their relationship to feedback during the bird’s own vocalization yet to be established. Despite these technical constraints, the robust convergence of our lesion, pharmacological, and electrophysiological data provides compelling evidence that HVC supplies the basal ganglia with the temporally precise sensory template required for vocal learning. Isolating the HVC(X) projection, recording the gate as it closes over learning, and testing how the gated tutor signal interacts with dopaminergic and cholinergic input at the cortico-striatal synapse identified by Schreiner et al. (2026) are natural next steps toward a complete circuit account of how a memorized model instructs vocal learning.

## Methods

### Animals and tutoring paradigm

All experiments were approved by the Regierung von Oberbayern (AZ ROB-55.2-2532.Vet_02-19-185, AZ ROB-55.2-2532.Vet_02-21-201). Beginning at ~30 dph, each juvenile was housed with a single live adult male tutor within a sound-isolation chamber, providing continuous social tutoring throughout the sensory and sensorimotor phases of song learning (Figure 1A).

### Bilateral Area X lesions

Bilateral lesions of Area X were performed at defined ages across the critical period. Birds were anesthetized with 1-3% isoflurane and placed in a stereotaxic apparatus with the head fixed at 20° beak-bar angle. This was necessary to bypass LMAN. Area X was targeted bilaterally using stereotaxic coordinates relative to the bifurcation of the midsagittal sinus: 5.8 AP, 1.5 ML, 2.1 DV (measured in mm). Electrolytic microlesions were made by passing 200 µA of current for 90 s through a tungsten electrode (A-M systems, 5 MΩ) at three penetrations per hemisphere (site 1: 5.8 A-P, 1.3 M-L, 2.6 mm deep, site 2: 5.8 A-P, 1.5 M-L, 2.8 mm deep, site 3: 5.8 A-P, 1.7 M-L, 2.6 mm deep). Control birds underwent identical handling but received no lesion (see surgery for intracellular recordings). In a subset of birds, a unilateral craniotomy was made over HVC for either subsequent pharmacological manipulation (0.3 to −0.3 A-P,1.9 to 2.6 M-L, head-angle: 65°) or for implanting a stimulating electrode (0.2 A-P, 2.3 M-L, 0.33 D-V) for orthodromic verification of X neurons receiving projections from HVC. After surgery, birds were returned to the tutor chamber to resume tutoring where developmentally appropriate. Precise quantification of lesion extent was limited by histological assessment being performed only in the adult brain, where compensatory reorganization may have altered the apparent lesion size, although the scar tissue remained reliably identifiable in all the birds. With this caveat, lesion extent was comparable across age groups (Fig S1) and therefore could not explain the observed differences in tutor song imitation.

### Sharp intracellular recordings

Area X was identified as described above. During the initial surgery, a small craniotomy was made over Area X and sealed with silicone elastomer (Kwik-Cast, World Precision Instruments) to allow later access. A custom-made horseshoe-shaped headplate was affixed to the skull using dental cement (Palladur) to facilitate recordings in awake, head-restrained animals.

On the day of recording, the headplate was secured to the recording apparatus, and the bird was stabilized in a custom-built restraining box. The silicone elastomer and dura were removed to expose the brain. Sharp intracellular recording electrodes were pulled immediately before use from borosilicate glass capillaries with filament (outer diameter, 1.0 mm; inner diameter, 0.5 mm) using a horizontal micropipette puller (P-97; Sutter Instrument Company). Electrodes were back-filled with 3 M potassium acetate and had impedances >100 MΩ.

To prevent desiccation of the exposed brain, a well was constructed around the craniotomy using silicone elastomer and filled with phosphate-buffered saline (PBS). Recording electrodes were advanced in 5-10 µm steps using a micromanipulator (MP-285; Sutter Instrument Company). Neurons encountered at depths of 2.1–3.8 mm with stable resting membrane potentials were targeted for recording. Membrane penetration was achieved using a brief depolarizing pulse. Following penetration, some neurons required transient hyperpolarizing pulse to stabilize the recording. The holding current was gradually removed, and all data were collected at 0 holding current.

Playback of the tutor song began once a stable recording had been established. Song presentations were delivered at randomized intervals exceeding 1.5 s. Neural and acoustic signals were acquired and analyzed offline using custom MATLAB software. Only neurons that remained stable for at least nine song presentations were included in the final analysis.

In 5 of 19 birds, neurons in Area X were confirmed by orthodromic identification following implantation of a stimulating electrode in HVC during the initial surgery.

### Pharmacological manipulation

In a separate cohort of birds (gabazine, *n* = 5, 66-89 dph; muscimol, n = 4, 61-74dph), intracellular recordings from Area X neurons were performed as described above while pharmacologically manipulating HVC. Gabazine (0.1 mM) or muscimol (5 mM) was applied topically to HVC using an absorbable gelatin sponge (SMI-SPON, SMI AG), as previously described in Vallentin et al. (2016). Control experiments were performed using PBS applied in the same manner.

### Histology and lesion verification

At the end of the experiment (≥90 dph), birds were deeply anesthetized and perfused with PBS followed by 4% paraformaldehyde. Post-fixation (with 4% PFA and 30% sucrose solution), brains were sectioned in a cryostat (Leica systems) into 50 µm thick sections and stained with 1% Cresyl Violet solution to confirm the location and extent of bilateral Area X damage. Only birds with confirmed electrode tracts in Area X, with minimal encroachment on surrounding striatum or overlying nuclei, were included in the analyses (examples shown in S2).

### Song recording

Song was recorded continuously in sound-isolation chambers using SAP 2011 ^39^ sampled at 44.1 kHz. Undirected song was captured across development, including immediately before and after each lesion, and at song crystallization (≥90 dph). Recordings were segmented into song bouts and motifs using amplitude-based segmentation in custom MATLAB software. For each bird, at least 30 representative crystallized adult motifs (>90 dph) was used for the tutor-copying analyses, and the corresponding adult tutor’s motifs served as the reference model.

### Acoustic similarity analysis-pre and post lesions

To quantify how similar a bird’s crystallized song was to (1) its own other renditions and (2) the tutor model, we computed pairwise acoustic similarity between motifs (average number of tutor motifs: 86 (range: 69-118 motifs); average number of tutee motifs: 97 (range: 36-140 motifs) using a sliding-window spectral cross-correlation approach ^40^. Briefly, each motif was converted to a spectrogram, similarity between two motifs was computed by cross-correlating their spectral content within a sliding temporal window and taking the peak correlation to align the two renditions in time, yielding a similarity score bounded between 0 and 1.

Two quantities were derived from these pairwise scores:

- *Self-similarity (self-stereotypy):* the mean similarity across all pairwise comparisons between distinct crystallized motifs of the same bird, indexing rendition-to-rendition consistency.
- *Tutor similarity*: the mean similarity across all pairwise comparisons between the tutee’s crystallized motifs and the tutor’s motifs, averaged over all tutor–tutee motif pairs, indexing how reliable the bird copied the model.

To capture overall imitation success in a single measure, we defined a *maturity score* as the product of self-stereotypy and tutor similarity (maturity = self-stereotypy × similarity to tutor) normalized by the similarity of the tutor to itself (tutor self-similarity). This score is high only when a bird sings a consistent song (high self-stereotypy) that also resembles the tutor (high tutor similarity).

### Acoustic feature extraction and dimensionality reduction

To visualize the acoustic relationship between tutor and the ultimately acquired tutee repertoire syllables, syllable waveforms were extracted from manually annotated syllables (tutee (average number of syllables per day): 226 syllables, (range: 39-446 syllables); tutor 1: 299 syllables, tutor 2: 180 syllables) in custom MATLAB software. For each syllable, a waveform segment extending 20 ms before syllable onset and 20 ms after syllable offset was extracted. To minimize differences in recording amplitude across files, each recording was normalized to a target root-mean-square (RMS) amplitude of 0.1. If normalization resulted in signal clipping, the waveform was rescaled to keep all sample values within the range of ±1.

Waveform segments were zero-padded to a common length, producing fixed-length vectors suitable for comparison across syllables. The resulting dataset was globally normalized by dividing all samples by the maximum absolute amplitude across the dataset.

Dimensionality reduction was performed using Uniform Manifold Approximation and Projection ^41^ implemented in MATLAB (10 output dimensions, n = 10 nearest neighbors, minimum distance = 0.05-0.005, correlation/cosine distance metric). The resulting 10-dimensional UMAP embeddings were subsequently subjected to principal component analysis (PCA). The first three principal components, retaining the vast majority of variance in the UMAP embedding, were used for subsequent HDBSCAN clustering ^42^. To identify suitable clustering parameters, HDBSCAN was evaluated across multiple combinations of minimum cluster size (5, 10, 20, 50, and minimum samples (default, 5, 10, and 20). For each parameter combination, clustering performance was quantified using the number of identified clusters, the proportion of syllables classified as noise, the mean silhouette coefficient (excluding noise points), the mean cluster persistence reported by HDBSCAN, and the median cluster membership probability. The parameter combination yielding the highest cluster stability while maintaining high membership probabilities and biologically interpretable clusters was selected for subsequent analyses. Syllables belonging to the same cluster were labelled as “learned” otherwise “not-learned”.

Before computing similarity scores, low-amplitude portions from each annotated syllable were trimmed using a 5-ms moving-average amplitude envelope and a threshold set to 5% of the maximum envelope value. Spectrograms were constructed using a 1024-sample Hamming window, 1000-sample overlap, and 1024-point FFT at 44.1 kHz and restricted to 500–7500 Hz, and linearly interpolated along the time dimension to 100 bins.

If any given cluster, contained only a single tutor and tutee syllable-type, the similarity score between the syllables was computed by calculating a median pairwise correlation on the 2D pre-processed spectrograms (as described above). For clusters where multiple tutee syllable-types overlapped with a tutor syllable, both were assumed to map to the same tutor syllable and a similarity score was computed by averaging scores from the comparison of individual tutee syllables with the tutor syllables. In case, none of the tutee syllables overlapped with any tutor syllable (not learned syllables), a similarity score was calculated by averaging all scores resulting from rendition-by-rendition comparison of this tutor syllable with all tutee syllables. Scores to individual elements in the birds own song were computed in a similar manner where if a given syllable in BOS did not have an analogue in the tutor song, it was evaluated against all tutor syllables and averaged. If song recordings were unavailable from the neural recording day, we used recordings from the nearest earlier day.

### Quantification and statistical analysis

Statistical details for each comparison, including the test used, the number of birds, and the resulting statistics, are reported in the corresponding figure legend and results. All analyses were performed in MATLAB (MathWorks) and no statistical methods were used to predetermine sample size.

### Spike waveform extraction and clustering

Action potentials were extracted from intracellular voltage recordings sampled at 40 kHz. Individual voltage traces were baseline corrected by subtracting a smoothed version of the signal. Spikes were detected in the playback window by setting a user-defined threshold manually by observing the neuron-specific amplitude. For each detected spike, a 6 ms waveform window (2 ms preceding and 4 ms following the spike peak) was extracted. Only complete waveforms were retained for subsequent analyses. Waveforms were normalized by dividing each waveform by its peak amplitude to minimize variability due to spike size while preserving waveform shape.

Dimensionality reduction was first performed using principal component analysis (PCA) on the normalized spike waveforms. The first 18 principal components were retained and standardized by dividing each component by its standard deviation. The resulting feature matrix was embedded into two dimensions using Uniform Manifold Approximation and Projection (UMAP) (nearest neighbors = 10, min dist = 0.05, ‘cosine distance’).

Spike waveforms were assigned to clusters using HDBSCAN (implemented in Python) applied to the UMAP embedding (as described above). Classification accuracy was ass-essed by calculating the proportion of neurons for which more than 80% of spikes were assigned to a single cluster.

Neurons that were not identified as spiking during the primary analysis window were reanalyzed using the entire recording duration to identify any spikes that may have occurred outside the initial analysis period. Neurons in which no spikes were detected throughout the complete recording were classified as non-spiking.

### Spike rate cross correlations

Raw voltage traces were analyzed offline using custom MATLAB scripts. Recordings were sampled at 40 kHz. For each trial, the trace was high-pass filtered by subtracting a smoothed version of the signal from the original trace. Putative spikes were detected during the playback and baseline windows separately and were used to generate a sliding-window PSTH (10 ms bins, 1 ms jumps). To determine whether stimulus-evoked activity significantly deviated from baseline firing variability, trial-shuffled surrogate PSTHs (n = 1000 shuffles) were generated. Significant increases and decreases in firing activity during playback were identified by comparing the actual PSTH to the 99th percentile of the shuffled distribution at each time point. Baseline activity was assessed similarly using the 99th percentile of the baseline shuffled distribution.

### Sliding window subthreshold correlations

Each neural trace was denoised to remove 50 Hz electrical noise and demeaned by subtracting the average membrane potential. Spikes from the resulting traces were removed using a threshold-based interpolation method. Membrane potential changes were quantified in a 30 ms sliding-window with advancements of 1 ms steps to capture the slow timescale of subthreshold dynamics. Every 30 ms bin was estimated as its trial-averaged membrane potential yielding a continuous average membrane potential trace with a resolution of 1ms. For each 30 ms window, zero-lag cross-correlations were calculated between all pairwise combinations of trials. The resulting correlation values were averaged across all trial pairs to obtain a mean subthreshold correlation value for each time point.

Stimulus-related changes in membrane potential and trial-to-trial correlation were identified relative to baseline (750 ms before playback onset) variability. For each measure, the baseline mean and standard deviation were calculated, and thresholds were defined as three standard deviations above and below the baseline mean. Epochs were detected by identifying contiguous periods during which both the membrane potential and correlation measure (as calculated above) exceeded these respective baseline thresholds. It was necessary to take the overlap of the two measures to avoid false positives possibly resulting from either a high cross-correlation between unmodulated trials or from jumps in membrane potential resulting from spiking activity.

Epochs during which the averaged membrane potential exceeded the upper threshold were classified as depolarizing response epochs, whereas periods falling below the lower threshold were classified as hyperpolarizing response epochs.

### Linear mixed effect models

Because multiple neurons were recorded from the same birds, and birds could share tutors and syllable identities, observations were not independent. We therefore used linear mixed-effects models to account for this hierarchical structure (Table S1):

Subthreshold response~Similarity+Age+Similarity*condition/playback+(1|random effects)

The subthreshold response (depolarization or hyperpolarization) was the outcome variable. Fixed effects included age at recording and the similarity of each juvenile syllable to its corresponding tutor syllable (for BOS Playback) or similarity of each tutor syllable to its corresponding juvenile syllable (for tutor playback). Where relevant, models also included the interaction between similarity and playback type (tutor song or bird’s own song [BOS]) or between similarity and treatment condition (gabazine or control).

We included random intercepts for tutor identity, syllable identity, bird identity, and cell identity, as appropriate for each model. Fixed-effect estimates and interaction terms were evaluated using a significance threshold of *p* < 0.05.

---

**Table S1:**
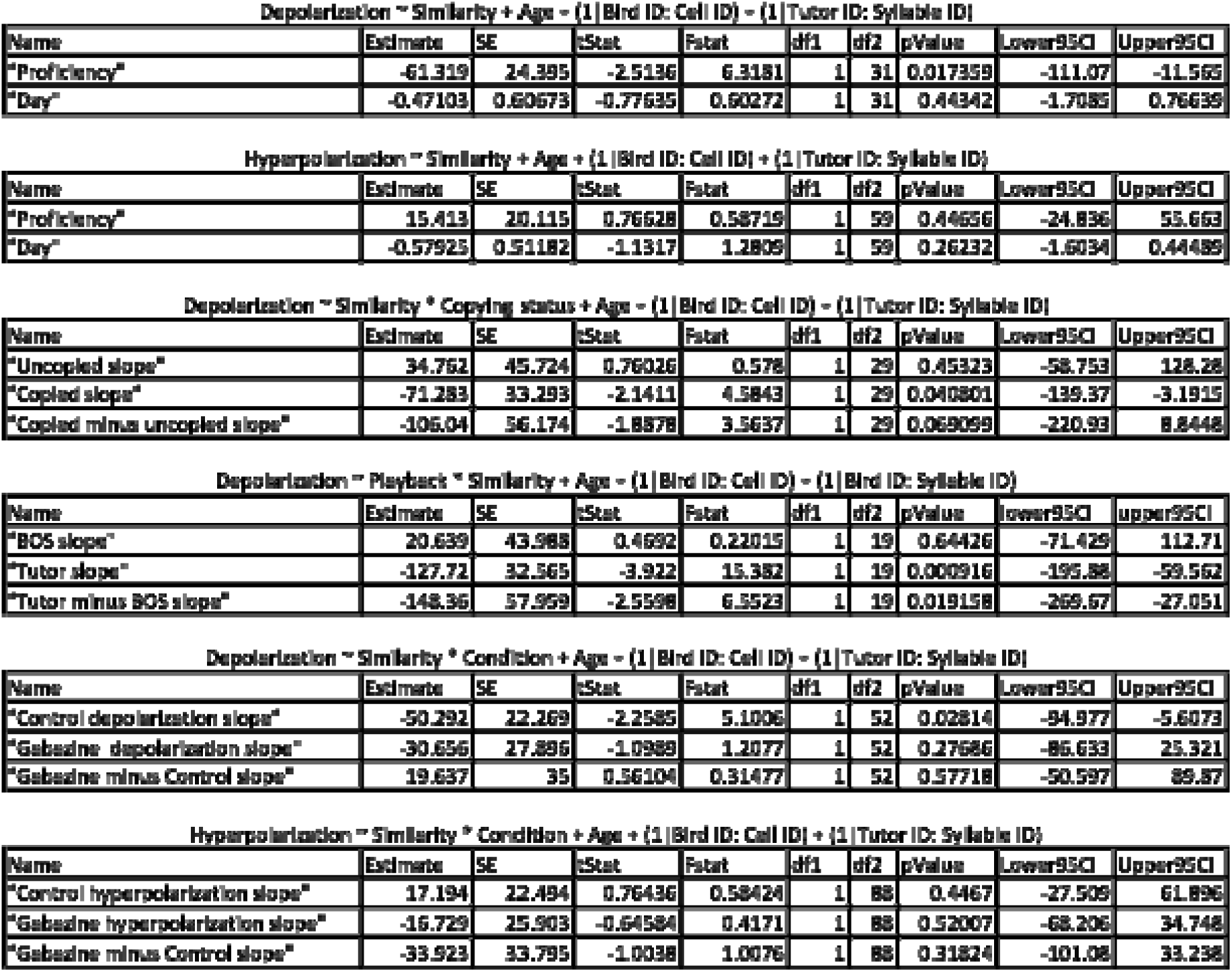
Linear mixed effect model results.

| Depolarization ~ Similarity + Age + (1\|Bird ID: Cell ID) + (1\|Tutor ID: Syllable ID) |  |  |  |  |  |  |  |  |  |
|---|---|---|---|---|---|---|---|---|---|
| Name | Estimate | SE | tStat | Fstat | df1 | df2 | pValue | Lower95CI | Upper95CI |
| "Proficiency" | -61.319 | 24.395 | -2.5136 | 6.3181 | 1 | 31 | 0.017359 | -111.07 | -11.565 |
| "Day" | -0.47103 | 0.60673 | -0.77635 | 0.60272 | 1 | 31 | 0.44342 | -1.7085 | 0.76639 |
| Hyperpolarization ~ Similarity + Age + (1\|Bird ID: Cell ID) + (1\|Tutor ID: Syllable ID) |  |  |  |  |  |  |  |  |  |
| Name | Estimate | SE | tStat | Fstat | df1 | df2 | pValue | Lower95CI | Upper95CI |
| "Proficiency" | 15.413 | 20.115 | 0.76628 | 0.58719 | 1 | 59 | 0.44656 | -24.836 | 55.663 |
| "Day" | -0.57925 | 0.51182 | -1.1317 | 1.2809 | 1 | 59 | 0.26232 | -1.6034 | 0.44489 |
| Depolarization ~ Similarity * Copying status + Age + (1\|Bird ID: Cell ID) + (1\|Tutor ID: Syllable ID) |  |  |  |  |  |  |  |  |  |
| Name | Estimate | SE | tStat | Fstat | df1 | df2 | pValue | Lower95CI | Upper95CI |
| "Uncopied slope" | 34.762 | 45.724 | 0.76026 | 0.578 | 1 | 29 | 0.45323 | -58.753 | 128.28 |
| "Copied slope" | -71.283 | 33.293 | -2.1411 | 4.5843 | 1 | 29 | 0.040801 | -139.37 | -3.1915 |
| "Copied minus uncopied slope" | -106.04 | 56.174 | -1.8878 | 3.5637 | 1 | 29 | 0.069099 | -220.93 | 8.8448 |
| Depolarization ~ Playback * Similarity + Age + (1\|Bird ID: Cell ID) + (1\|Bird ID: Syllable ID) |  |  |  |  |  |  |  |  |  |
| Name | Estimate | SE | tStat | Fstat | df1 | df2 | pValue | lower95CI | upper95CI |
| "BOS slope" | 20.639 | 43.988 | 0.4692 | 0.22015 | 1 | 19 | 0.64426 | -71.429 | 112.71 |
| "Tutor slope" | -127.72 | 32.565 | -3.922 | 15.382 | 1 | 19 | 0.000916 | -195.88 | -59.562 |
| "Tutor minus BOS slope" | -148.36 | 57.959 | -2.5598 | 6.5523 | 1 | 19 | 0.019158 | -269.67 | -27.051 |
| Depolarization ~ Similarity * Condition + Age + (1\|Bird ID: Cell ID) + (1\|Tutor ID: Syllable ID) |  |  |  |  |  |  |  |  |  |
| Name | Estimate | SE | tStat | Fstat | df1 | df2 | pValue | Lower95CI | Upper95CI |
| "Control depolarization slope" | -50.292 | 22.269 | -2.2585 | 5.1006 | 1 | 52 | 0.02814 | -94.977 | -5.6073 |
| "Gabazine depolarization slope" | -30.656 | 27.896 | -1.0989 | 1.2077 | 1 | 52 | 0.27686 | -86.633 | 25.321 |
| "Gabazine minus Control slope" | 19.637 | 35 | 0.56104 | 0.31477 | 1 | 52 | 0.57718 | -50.597 | 89.87 |
| Hyperpolarization ~ Similarity * Condition + Age + (1\|Bird ID: Cell ID) + (1\|Tutor ID: Syllable ID) |  |  |  |  |  |  |  |  |  |
| Name | Estimate | SE | tStat | Fstat | df1 | df2 | pValue | Lower95CI | Upper95CI |
| "Control hyperpolarization slope" | 17.194 | 22.494 | 0.76436 | 0.58424 | 1 | 88 | 0.4467 | -27.509 | 61.896 |
| "Gabazine hyperpolarization slope" | -16.729 | 25.903 | -0.64584 | 0.4171 | 1 | 88 | 0.52007 | -68.206 | 34.748 |
| "Gabazine minus Control slope" | -33.923 | 33.795 | -1.0038 | 1.0076 | 1 | 88 | 0.31824 | -101.08 | 33.236 |

## Acknowledgements

We thank members of the Vallentin lab for providing feedback on the initial versions of the manuscript; Animal care-takers at Max Planck Institute for Biological Intelligence for maintenance of our zebra finch colonies

## Funding

HORIZON EUROPE European Research Council (ERC)-2017-StG-757459 MIDNIGHT (DV)

Deutsche Forschungsgemeinschaft 327654276 - SFB 1315 (DV)

Deutsche Forschungsgemeinschaft 532521431 - FOR 5768 (DV)

Deutsche Forschungsgemeinschaft 547921981 - VA 742/6-1 (DV)

## Author contributions

Conceptualization: AA, DV

Methodology: AA, DV

Investigation: AA, DV

Visualization: AA, DV

Funding acquisition: AA, CS, DV

Project administration: AA, CS, DV

Supervision: DV

Writing – original draft: AA, DV

Writing – review & editing: AA, CS, DV

## Data and materials availability

Code and data for this project is available at https://github.com/vallentinlab

